# BRCA2 reversion mutation-independent resistance to PARP inhibition in prostate cancer through loss of function perturbations in the DNA pre-replication complex

**DOI:** 10.1101/2025.04.20.649668

**Authors:** Kyrie Pappas, Matteo Ferrari, Perianne Smith, Subhiksha Nandakumar, Zahra Khan, Serina B. Young, Justin LaClair, Marco Vincenzo Russo, Emmet Huang-Hobbs, Nikolaus Schultz, Wassim Abida, Wouter Karthaus, Maria Jasin, Charles Sawyers

## Abstract

Recent approvals of PARP inhibitors (PARPi) for BRCA-mutant metastatic castration resistant prostate cancer (mCRPC) necessitate an understanding of the factors that shape sensitivity and resistance. Reversion mutations that restore homologous recombination (HR) repair are detected in ∼50-80% of BRCA-mutant patients who respond but subsequently relapse, but there is currently little insight into why only ∼50% of BRCA-mutant patients display upfront resistance. To address this question, we performed a genome-wide CRISPR screen to identify genomic determinants of PARPi resistance in murine *Brca2*^Δ*/*Δ^ prostate organoids genetically engineered in a manner that precludes the development of reversion mutations. Remarkably, we recovered multiple independent sgRNAs targeting three different members (*Cdt1, Cdc6, Dbf4*) of the DNA pre-replication complex (pre-RC), each of which independently conferred resistance to olaparib and the next generation PARP-1 selective inhibitor AZD5305. Moreover, sensitivity to PARP inhibition was restored in *Brca2*^Δ*/*Δ^, Cdc6-depleted prostate cells by knockdown of geminin, a negative regulator of Cdt1, further implicating the critical role of a functional pre-RC complex in PARPi sensitivity. Furthermore, ∼50% of CRPC tumors have copy number loss of pre-RC complex genes, particularly *CDT1*. Mechanistically, prostate cells with impaired pre-RC activity displayed rapid resolution of olaparib-induced DNA damage as well as protection from replication fork degradation caused by Brca2 loss, providing insight into how Brca2-mutant cancer cells can escape cell death from replication stress induced by PARP inhibition in the absence of HR repair. Of note, a pharmacologic inhibitor that targets the CDT1/geminin complex (AF615) restored sensitivity to AZD5305, providing a potential translational avenue to enhance sensitivity to PARP inhibition in BRCA-mutant cancers.

**Significance:** Here, we address a major limitation in the effectiveness of PARP inhibitors in BRCA-mutant prostate cancer treatment: only ∼50% of patients respond despite clear genomic evidence of defective homologous recombination. Prior efforts to study PARP inhibitor resistance in prostate cancer have been plagued by the lack of suitable cell lines. We overcame this challenge using primary prostate organoids coupled with genome-wide CRISPR screening. The key finding is that loss of function mutations in the DNA pre-replication complex confer PARP inhibitor resistance. These genes map to chromosomal regions frequently lost in prostate cancer and could therefore serve as potential biomarkers of treatment response. Pharmacologic inhibition of geminin, a negative regulator of the pre-replication complex, can restore PARP inhibitor sensitivity.

## Introduction

Inhibitors of poly(ADP-ribose) polymerase (PARP) have transformed the treatment of patients with BRCA-mutant breast and ovarian cancers, which typically develop in a background of germline mutations in *BRCA1* or *BRCA2*. More recently PARPi have become a critical addition to the treatment of metastatic castration resistant prostate cancer (CRPC) where, unlike breast and ovarian cancer, ∼50% of BRCA mutations occur in the absence of an inherited germline mutation. Furthermore, *BRCA2* mutation/deletion is far more common in prostate cancer than BRCA1 loss of function, whereas in breast cancer the relative frequencies on *BRCA1* versus *BRCA2* mutation are nearly equal (1). Clinical responses to PARPi are seen in ∼50% of BRCA-mutant CRPC patients, as well as in patients with less common alterations that result in HR defects such as *PALB2, FANCA, BRIP1*, and *RAD51B*. Interestingly, response rates in CRPC patients whose tumors have mutations in other DNA damage repair (DDR) pathway genes such as *ATM*, *CDK12* and *CHEK2* are substantially lower, suggesting that many of these alterations may not result in HR defects of sufficient magnitude to confer synthetic sensitivity to PARP inhibition (2) (3).

Through its poly(ADP-ribose) parylation (PAR) activity, PARP1 modifies thousands of proteins involved in DNA damage response and plays a critical role in base excision repair, a key pathway in the repair of single-strand DNA breaks (4). In this context, the cytotoxicity of PARPi in BRCA-mutant cancers is explained by the inability of HR-deficient cells to repair the double strand DNA breaks that develop following PARP inhibition (5–7). Growing evidence suggests that the cytotoxicity of PARPi in HR deficient cells is not a consequence of blocking PARP enzymatic activity but rather a consequence of trapping PARP on DNA. Trapped PARP1-DNA complexes interfere with DNA replication, and deficiencies in replication fork protection alone (in the setting of intact HR) can result in synthetic lethality to PARPi in specific contexts (8). Indeed, the relative DNA trapping activity of next generation PARPi is more closely correlated with antitumor activity than with PARP enzyme inhibition (9, 10). Furthermore, genetic deletion or mutation of PARP1 confers resistance to PARPi therapy in BRCA-mutant cells, indicating that presence of the protein (perhaps for DNA trapping) rather than enzymatic activity is required for cytotoxic activity (11, 12). These insights, together with the efforts to improve the side effect profile of the first generation dual PARP1/2 inhibitor olaparib, have resulted in the development of several PARP1 selective compounds currently in clinical development (13) (14) (https://ascopubs.org/doi/10.1200/JCO.2024.42.16_suppl.TPS5123).

Despite the clincal success of PARPi across these different cancer settings, both upfront and acquired resistance pose significant limitations to achieving broader clinical benefit. In the case of acquired resistance, reversion mutations in BRCA1 or BRCA2 that restore HR proficiency are found in 50% or more of patients, including prostate cancer (15, 16). Genetic screens in BRCA-mutant cell lines and tumors have identified several mechanisms of reversion-independent resistance such as restoration of HR or replication fork protection, loss of PARP1 negative regulator PARG, alterations in PARP1, various epigenetic modifications, and PGP-mediated drug efflux that allow BRCA-mutant cells to survive despite being HR deficient, but the clinical importance of these mechanism are not yet clear (17–22). In prostate cancer, PARPi screens have been performed but not in HR-deficient settings, which poses challenges in knowing whether candidate resistance genes identified in this context will be informative in the clinical context where HR defects are predictive of clinical benefit (23, 24). For example, prior studies in LNCaP sublines with engineered BRCA2 deletion failed to acquire sensitivity to concentrations of PARP inhibitors used clinically (1). PARPi resistance studies in prostate cancer are further challenged by the paucity of PARPi-sensitive BRCA-mutant human prostate cancer cell lines that could be deployed for genetic screens without developing reversion mutations as the primary resistance mechanism (25–28).

To address this challenge we took advantage of recent developments in prostate organoid technology that allow rapid and efficient generation of complex genotypes that reflect those seen in patients (29, 30). Using this approach, we generated *Brca2*^Δ*/*Δ^ (31) primary mouse prostate organoids in three genetic contexts (*Brca2*^Δ*/*Δ^*Trp53*^Δ*/*Δ^; *Brca2*^Δ*/*Δ^*Trp53*^Δ*/*Δ^*Pten*^Δ*/*Δ^ and *Brca2*^Δ*/*Δ^*sgRb1*), documented acquisition of exquisite PARPi sensitivity, then performed genome-wide CRISPR screens across all three genotypes to identify sgRNAs enriched in the setting of PARPi treatment. Across all three genetic contexts, we recovered sgRNAs targeting *Cdt1*, *Cdc6* and *Dbf4* in various combinations, leading us to further investigate the role of the DNA pre-replication complex (pre-RC) in response to PARPi therapy. Here we show that loss of function perturbations in the pre-RC complex result in rapid resolution of DNA damage induced by PARP inhibition and rescue from defects in fork protection conferred by *Brca2* loss, ultimately allowing cells to become tolerant to olaparib-induced replication stress. Furthermore, genetic or pharmacologic inhibition of geminin, an upstream suppressor of pre-RC complex activity, restores sensitivity to PARPi in *Brca2*^Δ*/*Δ^ organoids with impaired pre-RC activity, providing a potential translational opportunity.

## Results

### Bi-allelic *Brca2* loss in murine prostate organoids results in orders of magnitude increased sensitivity to PARP inhibition

In the absence of HR-deficient human prostate cancer cell lines suitable for screening, we chose to model BRCA deficiency through Cre-mediated deletion in primary mouse prostate organoids freshly isolated from *Brca2^fl/fl^*mice (Brca2-null, deletion of N-terminal exons 3-4) (32, 33). In contrast to established prostate cancer cell lines, this approach provides the flexibility to model *Brca2* loss in primary cells together with co-occuring genomic alterations commonly seen in BRCA2-mutant human CRPC. Toward that end, we first generated prostate organoid cultures from mice containing a *Brca2^Fl/Fl^*allele alone or in combination with *Trp53^Fl/Fl^*and/or *Pten^Fl/Fl^* alleles. Ex vivo infection with Cre-expressing lentivirus virus was performed to generate organoid lines harboring homozygous loss of *Brca2* and *Trp53* ± *Pten* loss (*Brca2*^Δ*/*Δ^*Trp53*^Δ*/*Δ^ and *Brca2*^Δ*/*Δ^*Trp53*^Δ*/*Δ^ *Pten*^Δ*/*Δ^). Similarly, we also generated organoids with heterozygous *Brca2* loss (*Brca2*^+/Δ^ *Trp53*^Δ*/*Δ^ *Pten*^Δ*/*Δ^) to assess whether haploinsufficiency enhances sensitivity to PARP inhibition. We also modeled combined loss of *BRCA2* and *RB1* based on the fact that recurrent deletions of 13q23, spanning both the *BRCA2* and *RB1* gene loci, are often seen in CRPC (Fig 1A). *Rb1* was targeted in Cre-recombined *Brca2^Fl/Fl^ organoids* using sgRNAs (*Brca2*^Δ*/*Δ^ *sgRb1*), followed by ex vivo Cre recombination to delete *Brca2*. Deletion of *Trp53, Pten,* and *Rb1* in the respective organoid cultures was confirmed by western blot (Fig 1B). Recombination of the floxed *Brca2* locus was confirmed by PCR (Fig S1A-B). Histologic analysis across the *Brca2*-deleted organoid series revealed loss of the acinar/luminal/cystic features characteristic of wild-type organoids, likely a consequence of co-deletion of *Trp53*, *Pten* and *Rb1* as these same features are routinely observed following loss of these tumor suppressors in organoids with wildtype *Brca2* (Fig 1C).

**Figure 1:**
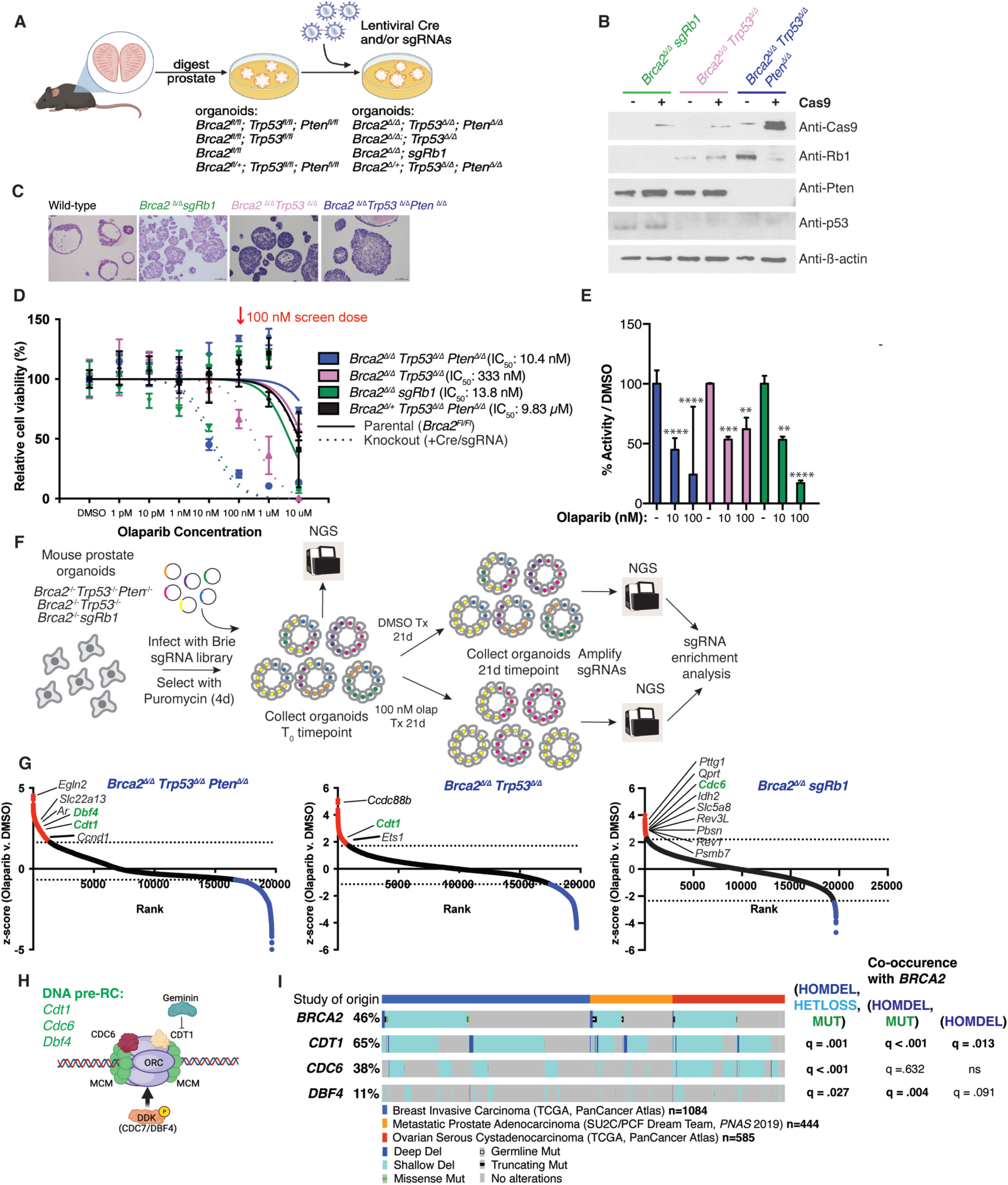
Genome-wide sgRNA screen in Brca2-deficient prostate organoids reveals loss of DNA pre-replication complex (Pre-RC) genes as a putative reversion-independent olaparib resistance mechanism. (A) Experimental flow for the generation of Brca2-deficient murine prostate organoid models. Prostates were harvested and genetic modifications were performed *in vitro,* as depicted. (B) Immunoblot confirming genotypes (*Rb1, Pten, Trp53*) of Brca2-deficient organoid models (*Brca2^Δ/Δ^ Trp53^Δ/Δ^ Pten^Δ/Δ^, Brca2^Δ/Δ^ Trp53^Δ/Δ^, Brca2^Δ/Δ^ sgRb1)* before and after introduction of stable Cas9 expression prior to screening. β-actin was used as loading control. See supplementary data for *Brca2* genotyping. (C) Representative images of H&E stains of *Brca2* wild-type and Brca2-deficient murine prostate organoid lines (D) 7-day olaparib dose response viability assay in Brca2-deficient murine prostate organoid series, where the genotypes are listed. Viability was assessed using the Incucyte and is displayed as percent inhibition relative to DMSO (vehicle) treatment. Error bars represent the mean ± SD of triplicate technical replicates. 100 nM dose used for screen is indicated (red arrow). Dose response curves were generated through nonlinear regression analysis. (E) PARP1 ELISA-based activity assay was used to quantify PARP1 activity in Brca2-deficient organoid lines (*Brca2^Δ/Δ^ Trp53^Δ/Δ^ Pten^Δ/Δ^, Brca2^Δ/Δ^ Trp53^Δ/Δ^, and Brca2^Δ/Δ^)* following 7-day treatment with DMSO (vehicle) or the indicated doses of olaparib. Data is displayed as % activity relative to DMSO (vehicle). Error bars represent the mean ± SD of triplicate technical replicates. Significance: Two-way ANOVA, Sidak’s correction, ****p<.0001, ***p<.001, **p<.01. (F) Experimental flow of genome-wide sgRNA screen (Brie lentiviral library) for olaparib resistance in Brca2-deficient murine prostate organoid series (*Brca2^Δ/Δ^ Trp53^Δ/Δ^ Pten^Δ/Δ^, Brca2^Δ/Δ^ Trp53^Δ/Δ^, and Brca2^Δ/Δ^ sgRb1* lines). Collection/sequencing timepoints are indicated (T_0_ is post-selection/pre-olaparib treatment, and DMSO /100 nM olaparib samples are collected after 21 days of treatment (∼10 cell doublings). (G) Rank ordered z-score plots (olaparib vs. DMSO treatment) from sgRNA screen for *Brca2^Δ/Δ^ Trp53^Δ/Δ^ Pten^Δ/Δ^, Brca2^Δ/Δ^ Trp53^Δ/Δ^, and Brca2^Δ/Δ^ sgRb1* lines (left to right). Each point represents one gene, and the dotted line represents the 5% FDR threshold for enrichment. Pre-RC genes are highlighted in green. Genes that pass the FDR threshold for enrichment are shown in red, and those that pass the threshold for depletion are shown in blue. (H) Diagram of Pre-RC complex, showing the screen hits *CDT1, CDC6*, and *DBF4* as subunits, among other proteins. (I) Oncoprint (generated by cBioPortal) showing homozygous deletions, shallow deletions, and mutations of *BRCA2, CDT1, CDC6*, and *DBF4*. Studies from breast, ovarian, and prostate cancer are included, and study details are shown. Oncoprint figure legend displays study of origin and the symbols for each type of alteration. Co-occurrence q-values are included for the indicated Pre-RC gene alterations (HOMDEL, HETLOSS, and/or MUT) with *BRCA2*.

Homozygous *Brca2* loss across all three genomic contexts (*Trp53*^Δ*/*Δ^; *Trp53*^Δ*/*Δ^*Pten*^Δ*/*Δ^; *sgRb1*) resulted in 30 to 1000-fold enhanced sensitivity to the PARPi olaparib, depending on the co-occurring genotype (with IC_50_s ranging from 10-333nM, versus 10uM in parental cells). This is notable because prior attempts to model the synthetic lethality to PARP inhibition seen with BRCA2 loss using human prostate cancer cell lines such as LNCaP yielded modest increases in olaparib sensitivity (low uM) that are not reflective of drug concentrations associated with clinical activity (23, 24). Importantly, mono-allelic loss of *Brca2* did not confer sensitivity to olaparib, with an IC_50_ of 10 uM, similar to that of the *Brca2^fl;fl^*lines prior to Cre recombination (Fig 1D). As expected, Parp1 activity (measured by ELISA) was inhibited by 50-80% at 10 and 100 nM olaparib concentrations in *Brca2*-deficient organoids (Fig 1E). Collectively, these data establish that murine prostate organoid cultures with bi-allelic loss of *Brca2* accurately mirror the synthetic lethality observed clinically in BRCA-mutant patients treated with PARPi. Furthermore, the large window in IC50 between sensitive and resistant genotypes provides an efficient experimental platform to identify genetic determinants of PARPi resistance through large scale CRISPR sceening. Finally, the specific *Brca2^fl/fl^* allele used to generate this model precludes the generation of *Brca2* reversion mutations, allowing us to focus the screen on identifying reversion mutant independent resistance mechanisms.

### Recovery of multiple components of the DNA pre-replication complex from genome-wide CRISPR screen for PARPi resistance

To perform the screen we introduced a Brie mouse lentiviral library with puromycin selection containing 4 sgRNAs per gene and 1000 non-targeting sgRNAs (34) into organoid cultures of three Brca2-deficient genotypes (*Brca2*^Δ*/*Δ^ *Trp53*^Δ*/*Δ^; *Brca2*^Δ*/*Δ^ *sgRb1*; *Brca2*^Δ*/*Δ^ *Trp53*^Δ*/*Δ^ *Pten*^Δ*/*Δ^) at a multiplicity of infection of 0.3, and screens were run in duplicate for each organoid line. Cultures were selected for 4 days post-infection, a portion was collected (T_0_), and cells were passaged for 21 days (roughly 10 doublings) in vehicle or 100 nM olaparib, then harvested for sgRNA amplification and next-generation sequencing (NGS) to identify sgRNAs selectively enriched by olaparib treatment (Fig 1F). Using an FDR enrichment threshold of 5% (experimentally determined for each organoid line based on distribution of the nontargeting sgRNAs) together with a requirement of at least three of four sgRNAs per gene for hit calling (see Materials and methods for details), we generated a list of genes selectively depleted following olaparib treatment in *Brca2*^Δ*/*Δ^ *Trp53*^Δ*/*Δ^ organoids (n= 487), *Brca2*^Δ*/*Δ^ *sgRb1* organoids (n=149) and *Brca2*^Δ*/*Δ^ *Trp53*^Δ*/*Δ^ *Pten*^Δ*/*Δ^ organoids (n=322) (Fig 1G, Spreadsheet S1).

We then used gene set enrichment analysis (GSEA) to identify pathways represented by these genes, which revealed striking enrichment of multiple gene sets involved in DNA replication and DNA damage response (DDR) (Fig S2). Through this lens, we then examined specific genes within these pathways and noted enrichment of multiple independent sgRNAs targeting three different members (*Cdt1, Cdc6, Dbf4*) of the DNA pre-replication complex (pre-RC), a multi-protein complex that controls the spacing and timing of replication fork origin licensing to ensure that DNA is replicated once per cell cycle (depicted in Fig 1H). Furthermore, enrichment of at least one of these three core pre-RC components was seen across all three Brca2-deficient organoid cultures (Fig 1G).

Because a primary goal of the screen was to identify genes whose mutation/deletion might mitigate response to PARPi therapy in CRPC patients, we examined publically available prostate cancer genomic datasets for copy number loss of *CDT1, CDC6* and *DBF4* and whether these losses overlap with *BRCA2* mutation status (35, 36). Of these, *CDT1* loss is most common (65% of cases have heterozygous or homozygous loss) and these losses overlap significantly with BRCA2 loss. *CDC6* and *DBF4* loss are much less common but also show significant overlap with BRCA2 loss when both heterozygous and homozygous loss are both considered (Fig 1I). Based on these potentially relevant clinical associations, we focused on all three pre-RC genes for deeper investigation.

### Resistance to PARP inhibition conferred by impaired pre-RC complex function is rescued by geminin knockdown

Loss of *PARP1* has been reported to promote PARPi resistance in multiple contexts, and Parp1 sgRNAs were recovered from the screen in *Brca2*^Δ*/*Δ^ *Trp53*^Δ*/*Δ^ organoids, likely implicating the DNA trapping mechanism for PARPi-induced synthetic lethality as opposed to inhibition of enzymatic PARylation activity (11). To validate PARP1 as a screen hit, we expressed two independent sgRNAs against *Parp1* in *Brca2*^Δ*/*Δ^ *sgRb1* and *Brca2*^Δ*/*Δ^ *Trp53*^Δ*/*Δ^ *Pten*^Δ*/*Δ^ organoids (Fig 2A-B). Organoids tolerated PARP1 knockout and were resistant to olaparib, presumably due to loss of DNA trapping.

**Figure 2:**
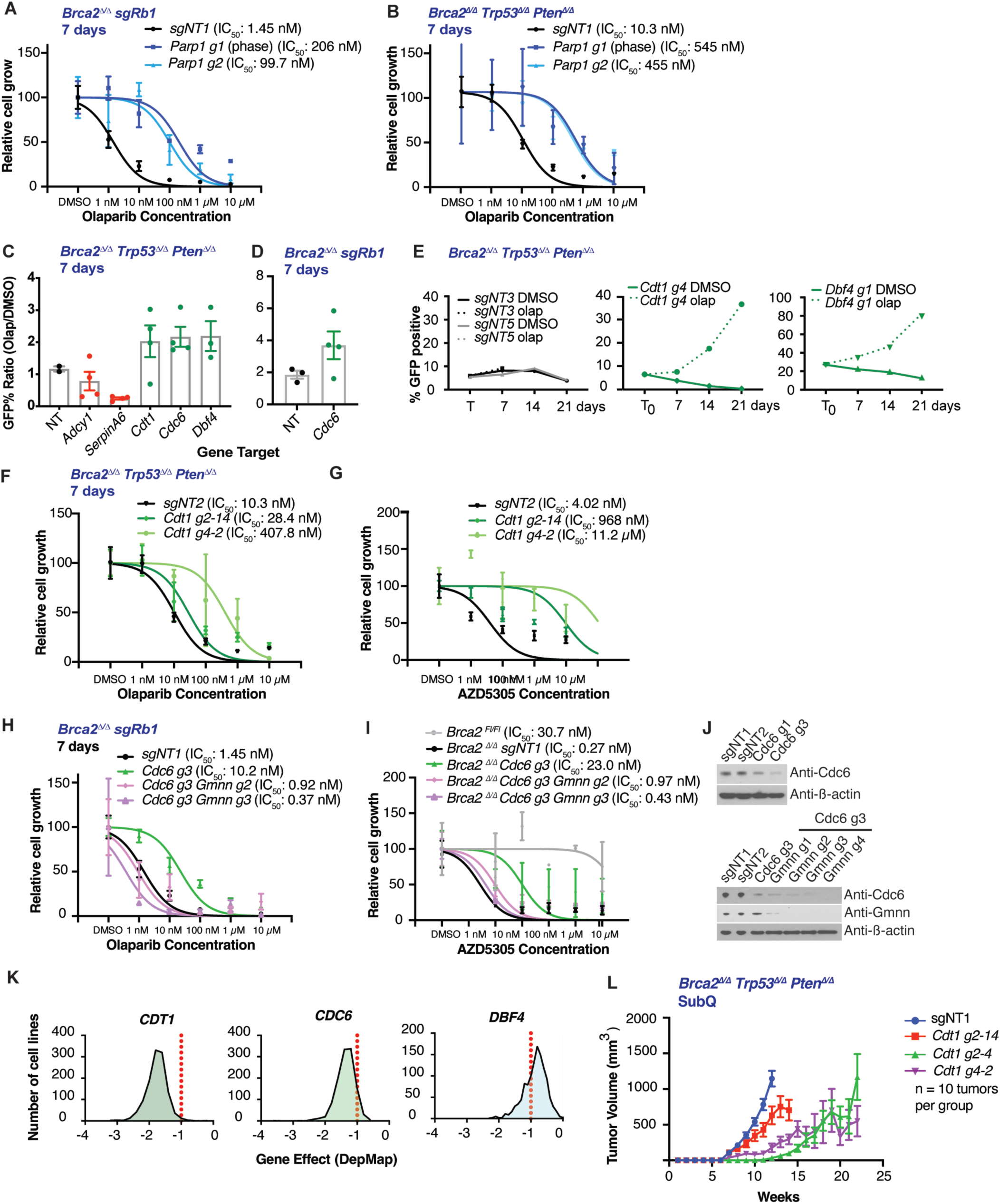
Loss of Pre-RC genes is tolerated and drives PARPi resistance in Brca2-deficient organoid models of mCRPC. 7-day olaparib dose response viability assay in (A) *Brca2^Δ/Δ^ Trp53^Δ/Δ^ Pten^Δ/Δ^, and (B) Brca2^Δ/Δ^ sgRb1* murine prostate organoids stably expressing the indicated sgRNAs (*sgNT1, Parp1 g1, Parp1 g2*). Viability was quantified using the Incucyte and is displayed as percent inhibition relative to DMSO (vehicle) treatment. Error bars represent the mean ± SD of triplicate technical replicates. Dose response curves were generated through nonlinear regression analysis. (C-D) GFP competition assays using indicated sgRNAs in LentiCRISPR-GFP treated with 100 nM olaparib or DMSO (vehicle) for 7 days in (C) *Brca2^Δ/Δ^ Trp53^Δ/Δ^ Pten^Δ/Δ^*, and (D) *Brca2^Δ/Δ^ sgRb1* organoids. Data is displayed as the ratio of %GFP in olaparib/DMSO. Error bars represent the mean ± SEM, and each point is an individual sgRNA. (E) Time course GFP competition assays for the indicated sgRNAs after 100 nM olaparib or DMSO (vehicle) treatment for 7, 14, and 21 days in *Brca2^Δ/Δ^ Trp53^Δ/Δ^ Pten^Δ/Δ^* organoids. Data is displayed as % GFP. Olaparib is the dashed line, DMSO (vehicle) is the solid line. GFP was quantified using flow cytometry for (C-E). (F-G) 7-day olaparib dose response viability assay in *Brca2^Δ/Δ^ Trp53^Δ/Δ^ Pten^Δ/Δ^* organoids stably expressing the indicated sgRNAs (*sgNT2, Cdt1 g2-14, Cdt1 g4-2*) in LentiCRISPR-GFP treated with (F) olaparib, or (G) AZD5305. Viability was quantified using the Incucyte and is displayed as percent inhibition relative to DMSO (vehicle) treatment. (F-G) Error bars represent the mean ± SD of triplicate technical replicates. Dose response curves were generated through nonlinear regression analysis. (H-I) 7-day olaparib dose response viability assay in *Brca2^Δ/Δ^ sgRb1* organoids stably expressing the indicated sgRNAs *(sgNT1, Cdc6 g3, Gmnn g2, Gmnn g3)* in LentiCRISPR-GFP treated with (H) olaparib, or (I) AZD5305. Viability was quantified using the Incucyte, and is displayed as percent inhibition relative to DMSO(vehicle) treatment. (H-I) Error bars represent the mean ± SD of triplicate technical replicates. Dose response curves were generated through nonlinear regression analysis. (J) Immunoblot assessing protein levels of Cdc6 and Geminin in *Brca2^Δ/Δ^ sgRb1* organoids stably expressing the indicated sgRNAs *(sgNT1, sgNT2, Cdc6 g1, Cdc6 g3, Gmnn g2, Gmnn g3)* in LentiCRISPR-GFP. β-actin was used as loading control. (K) Histograms of aggregate data from the Cancer Dependency Map (DepMap) displaying the effect on viability when expressing sgRNAs targeting *CDT1*, *CDC6*, and *DBF4* (left to right) in 1178 cell lines. The gene effect value of -1 is the standard cutoff for essential genes and is indicated by the dashed red line. (L) *Brca2^Δ/Δ^ Trp53^Δ/Δ^ Pten^Δ/Δ^* organoids expressing the indicated sgRNAs (*sgNT1, Cdt1 g2-14, Cdt1 g2-4, Cdt1 g4-2*) in LentiCRISPR-GFP were injected into NSG mice, tumor volume was measured weekly, displayed in mm^3^. Error bars represent mean ± SEM, n = 10 tumors per group.

To confirm the robustness of our sgRNA hit calling and validate the role of pre-RC complex in PARPi sensitivity, we used GFP competition assays to compare the relative enrichment of all pre-RC complex sgRNA hits versus various controls (non targeting sgRNAs as well as non-enriched sgRNAs) in *Brca2*^Δ*/*Δ^ *Trp53*^Δ*/*Δ^ *Pten*^Δ*/*Δ^ and *Brca2*^Δ*/*Δ^*sgRb1* organoids (see methods for details on enrichment screen). sgRNAs targeting pre-RC complex members *Cdt1, Cdc6* and *Dbf4* were enriched in 100 nM olaparib treatment versus DMSO (vehicle), whereas sgRNAs targeting *Adcy1* or *SerpinA6* (not enriched in the screen) and nontargeting sgRNAs (sgNT3, sgNT5) were not (Fig. 2C-E).

With the growing recognition that PARP1 is likely the critical target for PARPi-induced synthetic lethality in HR defective cancers, there is intense interest in developing compounds with improved PARP1 selectivity. AZD5305 (saruparib) is one such compound currently in phase 3 clinical development in CRPC (13) (14) (https://ascopubs.org/doi/10.1200/JCO.2024.42.16_suppl.TPS5123). Side-by-side viability assays comparing olaparib and AZD5303 confirmed multi-log enhanced sensitivity in *Brca2*^Δ*/*Δ^ *Trp53*^Δ*/*Δ^ *Pten*^Δ*/*Δ^ organoids as well as rescue by sgRNA targeting of *Cdt1*, a core component of the pre-RC (Fig 2F-G, Fig S3).

The pre-RC complex is tightly regulated, most notably by geminin which sequesters Cdt1 through physical interaction when origin licensing is not needed. Having validated loss of pre-RC complex as a cause of PARPi resistance, we next asked if we could restore PARPi sensitivity through geminin knockdown. Because the sequestering effect of geminin is specific to Cdt1, we first established PARPi resistance through introduction of sgRNAs targeting Cdc6 in *Brca2*^Δ*/*Δ^ *sgRb1* organoids, then similarly targeted geminin to release any sequestered Cdt1 with the goal of potentially restoring pre-RC activity. Remarkably, combined deletion of *Cdc6* and geminin restored sensitivity to olaparib and to AZD5305 to levels comparable to parental cells (Fig 2H-J). Collectively, these results confirm the importance of the pre-RC complex as a determinant of PARPi sensitivity.

It is worth noting that *CDT1*, *CDC6* and *DBF4* are all classified as essential genes in yeast and human in the most recent update of the DEG database (37). To better understand how *BRCA2*-mutant prostate organoids can tolerate deletion of these pre-RC complex genes, we examined the dependency profiles of *CDT1, CDC6* and *DBF4* in the cancer cell line dependency map (DepMap; https://depmap.org/portal/). Using a gene effect score of <-1 as a threshold for dependency, *CDT1* loss was not tolerated across most cancer cell lines whereas the consequences of *CDC6* and *DBF4* loss were less broad across the panel (Fig 2K). To investigate the consequences of Cdt1 loss in our prostate model, we evaluated the *in vivo* tumorigenicity of *Brca2*^Δ*/*Δ^ *Trp53*^Δ*/*Δ^ *Pten*^Δ*/*Δ^ organoids with and without *Cdt1* deletion. Three independent clonal *Cdt1*-edited sublines all gave rise to tumors, albeit with slower growth rates than parental Cdt1-intact organoids (Fig 2L). Importantly, complete *Cdt1* deletion was confirmed by isolating clones and sequencing across the sgRNA-edited locus in all three organoid sublines prior to injection, (clones characterized in Fig S3A-B. The resulting tumors also retained a high fraction of *Cdt1* edited alleles (Fig S4-S6), indicating that *Cdt1* loss is tolerated in this genomic context.

### PARPi-induced DNA damage is resolved and stalled replication forks recover in *Brca2*^Δ*/*Δ^ cells with impaired pre-RC function

PARPi induce DNA damage. In HR-competent cells this damage is repaired but, in cells lacking HR, dsDNA breaks accumulate resulting in cytotoxicity (7). To determine whether PARPi-induced DNA damage in *Brca2*^Δ*/*Δ^ prostate organoids with pre-RC complex dysfunction persists or is resolved, we measured γH2AX foci, a widely used marker of DNA damage, after 4 hours of olaparib treatment. As expected, γH2AX foci were induced in *Brca2*^Δ*/*Δ^ *sgRb1*and *Brca2*^Δ*/*Δ^ *Trp53*^Δ*/*Δ^ *Pten*^Δ*/*Δ^ organoids, indicative of the accumulation of DSBs (Fig 3A-B). In *Brca2*^Δ*/*Δ^ *sgRb1* organoids lacking Cdc6, these foci were resolved by 4 hours but not in those with concurrent geminin knockdown (Fig 3A). Similarly, γH2AX foci were resolved at 4 hours in *Brca2*^Δ*/*Δ^ *Trp53*^Δ*/*Δ^ *Pten*^Δ*/*Δ^ organoids with Cdt1 knockdown (Fig 3B). Taken together, these results establish that reversion mutation independent resistance to PARPi conferred by loss of pre-RC complex mutation is associated with resolution rather than tolerance of PARPi induced DNA damage.

**Figure 3:**
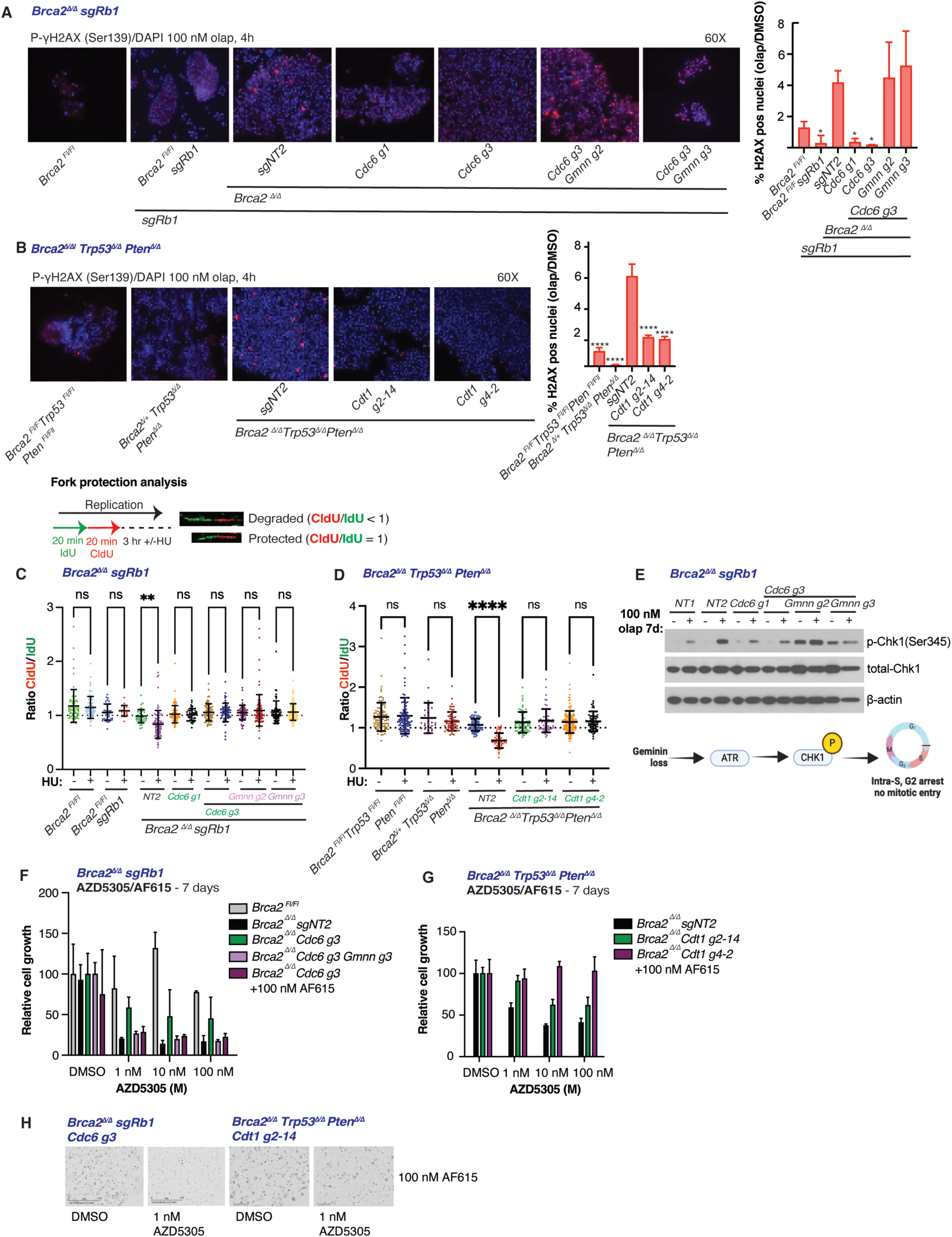
Loss of Pre-RC genes leads to PARPi resistance through the resolution of double strand breaks and rescued fork protection and can potentially be exploited by targeted therapy. (A-B) Representative 60X composite immunofluorescence images for γ-H2AX (red) and DAPI (blue) following 100 nM olaparib treatment for 4 hours, quantifications shown on right. (A) *Brca2^Δ/Δ^ sgRb1* organoids stably expressing the indicated sgRNAs *(sgNT1, sgNT2, Cdc6 g1, Cdc6 g3, Gmnn g2, Gmnn g3)* in LentiCRISPR-GFP. *Brca2^Fl/Fl^ and Brca2^Fl/Fl^sgRb1* organoids were included as controls. (B) *Brca2^Δ/Δ^ Trp53^Δ/Δ^ Pten^Δ/Δ^* organoids expressing the indicated sgRNAs (*sgNT1, Cdt1 g2-14, Cdt1 g4-2*) in LentiCRISPR-GFP. *Brca2^Fl/Fl^ Trp53^Fl/Fl^ Pten^Fl/Fl,^* and *Brca2^Δ/+^ Trp53^Δ/Δ^ Pten^Δ/Δ^* organoids were included as controls. Quantifications (A-B) are the ratio of % positive γ-H2AX in olaparib/DMSO treated organoids. Error bars represent the mean ± SD of triplicate technical replicates, and significance was calculated by 1-way ANOVA relative to the nontargeting (sensitive) organoid line. (C-D) DNA fiber analysis was performed where cells were treated with IdU (green) and CldU (red) for 20 minutes each, followed by hydroxyurea (HU) for 3h. Data is displayed as a ratio of IdU/CldU track length, where a ratio < 1 indicates a degraded fork, and a ratio of 1 indicates a protected fork. (C) DNA fiber analysis in *Brca2^Δ/Δ^ sgRb1* organoids stably expressing the indicated sgRNAs *(sgNT1, sgNT2, Cdc6 g1, Cdc6 g3, Gmnn g2, Gmnn g3)* in LentiCRISPR-GFP. *Brca2^Fl/Fl^ and Brca2^Fl/Fl^sgRb1* organoids were included as controls. (D) DNA fiber analysis in *Brca2^Δ/Δ^ Trp53^Δ/Δ^ Pten^Δ/Δ^* organoids expressing the indicated sgRNAs (*sgNT1, Cdt1 g2-14, Cdt1 g4-2*) in LentiCRISPR-GFP. *Brca2^Fl/Fl^ Trp53^Fl/Fl^ Pten^Fl/Fl,^* and *Brca2^Δ/+^ Trp53^Δ/Δ^ Pten^Δ/Δ^* organoids were included as controls. (E) Immunoblot assessing protein levels of p-Chk1(Ser345) and total Chk1 in *Brca2^Δ/Δ^ sgRb1* organoids expressing the indicated sgRNAs *(sgNT1, sgNT2, Cdc6 g1, Cdc6 g3, Gmnn g2, Gmnn g3)* in LentiCRISPR-GFP following DMSO (vehicle) or 100 nM olaparib treatment for 7 days. β-actin was used as loading control. (F) Viability assay in *Brca2^Δ/Δ^ sgRb1* organoids expressing the indicated sgRNAs (sgNT2, Cdc6 g3, Gmnn g3) in LentiCRISPR-GFP following treatment with DMSO (vehicle) or the indicated doses of AZD5305 and AF615 for 7 days. *Brca2^Fl/Fl^*organoids were included as controls. (G) Viability assay in *Brca2^Δ/Δ^ Trp53^Δ/Δ^ Pten^Δ/Δ^* organoids expressing the indicated sgRNAs (*sgNT1, Cdt1 g2-14*) in LentiCRISPR-GFP following treatment with DMSO (vehicle) or the indicated doses of AZD5305 and AF615 for 7 days. *Brca2^Fl/Fl^ Trp53^Fl/Fl^* were included as controls. Viability assay was quantified using the Incucyte and is displayed as percent inhibition relative to DMSO (vehicle) treatment. Error bars represent the mean ± SD of triplicate technical replicates. (H) Representative photos for indicated organoid lines following AZD5305/AF615 combination treatment. Photos were captured by the Incucyte. ****p<.0001, **p<.01, *p<.05.

In addition to its role in HR, one of the central functions of Brca2 in DNA replication is to protect stalled replication forks from collapse. Since pre-RC complex deficient *Brca2*^Δ*/*Δ^ organoids can overcome the genotoxic stress of PARP inhibition in the absence of HR repair, we asked whether fork protection was restored in resistance phenotypes driven by loss of pre-RC activity. We used DNA fiber analysis to visualize individual replication forks under hydroxyurea (HU)-mediated replication stress (Fig. 3C). *Brca2^Δ/Δ^sgRb1* murine prostate organoids displayed the expected fork protection defect caused by Brca2 deficiency (CldU/IdU ratio <1), which was not present in *Brca2^Fl/Fl^ or Brca2^Fl/Fl^sgRb1* organoids (Fig. 3C). Upon introduction of sgRNAs against Cdc6, the fork protection defect was fully rescued (Fig. 3C). Similarly, *Brca2^Δ/Δ^ Trp53^Δ/Δ^ Pten^Δ/Δ^* organoids displayed an identical fork protection defect, which was rescued by sgRNAs against *Cdt1* (Fig. 3D). As expected, fork speed was not affected by loss of Pre-RC members (Fig. S7A-B).

Interestingly, although the addition of Gmnn sgRNAs concurrently with Cdc6 loss restored PARPi sensitivity, the fork protection defect was not similarly rescued (Fig. 3C), suggesting that Geminin loss may promote sensitivity through an alternative mechanism. These results demonstrate that loss of pre-RC members can rescue the fork protection defect conferred by Brca2 loss and confer PARPi resistance. The finding that Geminin loss does not restore the fork protection defect yet confers PARPi sensitivity likely reflects the distinct roles of Cdc6 in origin licensing during G1, and Gmnn in regulating/sequestering Cdt1 during S and G2. Geminin loss has the potential to be more detrimental, resulting in origin licensing by Cdt1 at inappropriate times in the cell cyle, leading to mitotic catastrophe.

To further investigate how *Brca2*^Δ*/*Δ^ organoids respond to the DNA damage induced by PARPi, we interrogated the ATR/Chk1 checkpoint and found that *Brca2^Δ/Δ^sgRb1* organoids activate p-Chk1 in response to olaparib treatment in cells expressing control sgRNAs (NT1, NT2) and Cdc6 (Fig 3E, left panels). Of note, Gmnn knockdown on top of Cdc6 knockout resulted in constitutive Chk1 phosphorylation at baseline, in the absence of PARPi treatment (Fig. 3E), which increased further following olaparib exposure. This result is consistent with previous work showing that Geminin loss results in increased baseline activation of the intra-S checkpoint due to over-replication, ultimately leading to G2-arrest and mitotic catastrophe (38, 39). Whereas pre-RC dysfunction is sufficient to confer PARPi resistance in the absence of HR (in part through rescue of stalled replication forks), we posulate that geminin loss causes an increase in replication stress above the threshold for viability.

A recent report describing a small molecule inhibitor of the Geminin-CDT1 interaction (AF615) (40) prompted us to investigate the activity of this compound in *Brca2^Δ/Δ^ sgRb1 Cdc6 g3* organoids treated with escalating concentrations of AZD5305 (Fig. 3F, H). Resistance to AZD5305 driven by loss of Cdc6 was completely reversed by AF615, even at 1 nM AZD5305. Conversely, AF615 did not restore AZD5305 sensitivity to *Brca2^Δ/Δ^ Trp53^Δ/Δ^ Pten^Δ/Δ^ sgCdt1* organoids, an expected result because the sequestering effect of geminin requires Cdt1 (Fig. 3G, H). These results suggest a potential translational opportunity to explore the use of Geminin inhibitors to restore PARPi sensitivity in BRCA2-deficient cancers in certain contexts.

## Discussion

The treatment landscape of BRCA-mutant cancers has been revolutionized by PARPi therapy and continues to improve with the development of more potent, PARP1-selective compounds that enhance tolerability and should facilitate exploration of combination therapies. Nonetheless, PARPi resistance is a significant challenge that limits long term clinical benefit. In patients who relapse after an initial response to therapy, reversion mutations in the BRCA locus that restore HR repair are, by far, the most frequent cause of acquired resistance (15). However, our understanding of upfront resistance (i.e. failure to respond to initial treatment) in patients is less mature despite multiple efforts to define genetic determinants of PARPi sensitivity and resistance through CRISPR screens using experimental models.

Here we have approached this question through the lens of *BRCA*-mutant prostate cancer which, in contrast to breast and ovarian cancer, is largely a *BRCA2*-mutant disease with a higher percentage of somatic rather than germline cases, as well as frequent 13q copy number loss spanning the *BRCA2* gene. Prior PARPi screens in prostate cancer have been limited by a paucity of available BRCA-mutant human prostate cancer cell lines, as well as by the inability to accurately replicate the synthetic lethality phenotype of PARPi treatment using isogenic *BRCA*-wildtype/*BRCA*-mutant cell line models. Our approach differs in that we used primary mouse prostate organoid cultures to generate *Brca2*-mutant genotypes that, gratifyingly, mirror those seen in patients and acquire exquisite PARPi sensitivity (orders of magnitude reduction in IC_50_, low nanomolar range for olaparib). Furthermore, the genetic approach used to delete *Brca2* precludes the development of reversion mutations, allowing us to focus the screen on reversion-independent PARPi resistance mechanisms.

Among many validated hits, we were struck by the recovery of multiple independent sgRNAs to three core members of the DNA pre-RC (*Cdt1, Cdc6, Dbf4*) across all three of the genetic contexts used in the screen. We show that loss of these individual pre-RC members promotes resistance through resolution rather than tolerance of DNA damage and through rescued fork protection, allowing S-phase completion and cell cycle progression. Further efforts are needed to elucidate possible alternative DNA repair pathways facilitating the resolution of DNA damage following PARPi treatment in cells with impaired Pre-RC function and whether trapping of Parp1 by PARPi influcences these resistance phenotypes. In addition to γH2AX, measuring 53BP1 and RPA foci formation in resistant cells could provide more resolution on the type of DNA damage being resolved. Depletion of fork reversal enzymes has been attributed to increased fork protection, a finding that warrants futher investigation in the context of PARPi resistance conferred by pre-RC loss. The finding that enhancing pre-RC activity through *Gmnn* knockout in Cdc6-deficient organoids restores PARPi sensitivity not only validates that loss of pre-RC function confers PARPi resistance but also provides a potential path to address this challenge clinically through small molecule disruption of the Geminin-CDT1 complex (40). The Geminin knockout experiments also provide insight into the mechanism of restored PARPi sensitivity, likely through the higher level of baseline replication stress (as measured by elevated pChk1) that can no longer be tolerated following PARPi exposure.

The fact that genomic loss of several pre-RC complex proteins (CDT1, CDC6, DBF4) is evident in CRPC patients raises the possibility that these alterations could be genomic determinants of upfront PARPi response. Our preliminary efforts to address this possibility, while intriguing, are limited by small cohorts of genomically annotated patients with longitudinal clinical response data. As larger cohorts of CRPC patients treated with PARPi accrue, it will be important to specifically annotate copy number and mutation status of pre-RC complex genes as these are not routinely included in most cancer gene sequencing panels. Another important question is whether the pre-RC findings reported here in the BRCA2-mutant setting extend to cancers with BRCA1 loss, and whether the genetic background, such as loss of p53, PTEN, and/or RB1, influences the resistance phenotype. Loss of Cdt1 promoted resistance in both of the p53-deficient organoid lines tested in our study, though larger data cohorts are needed to fully understand the complex relationships between these resistance mechanisms and genetic background. It is also worth considering whether loss of function alterations in the pre-RC could be a more generalizable mechanism of resistance to other chemotherapies that induce replication stress such as topoisomerase inhibitors. Further efforts are also warranted to determine whether these resistance mechanisms are relevant to genetic backgrounds other than BRCA1/2 loss that result in HR and/or replication fork protection deficiencies.

## Materials and Methods

### Establishment of organoid cell lines

Prostates were harvested from a series of *Brca2*-floxed mouse strains: *Brca2^fl/fl^*, *Brca2^fl/fl^ Trp53 ^fl/fl^*, and *Brca ^fl/fl^ Trp53 ^fl/fl^ Pten ^fl/fl^*. The *Brca2^Fl/Fl^*allele used was previously published and does not produce functional protein (Brca2-null) (31). Prostates were minced, digested with 5 mg/mL collagenase type II followed by TrypLE (ThermoFisher, 12605010), and cultured according to published protocols (34, 41). These cultures were used to establish the Brca2-deficient organoid models for this study, and additional detail is provided below.

### Organoid cell culture

Organoid lines were cultured at at 37°C, 5% CO_2_ in Matrigel (Corning, 356231), where 35 uL domes of matrigel containing the cells/organoids are placed on standard untreated plates for nonadherent cells (any brand/size), allowed to solidify for 20 min, and media is placed on top to fully submerge the Matrigel discs. Mouse prostate organoids were grown in standard mouse prostate organoid media composed of 1X Advanced DMEM/F-12 (Thermo Fisher Scientific, 12634028), 10% Noggin conditioned media, 5% R-Spondin conditioned media, 1% Pen/strep (Fisher Scientific, 15-140-122), 1% Glutamax (Thermo Fisher Scientific, 35050079), 10 mM HEPES (pH 7.4), 1X B27 (Fisher Scientific, 17504-044), 200 nM A83-01 (Tocris, 2939), and 1.25 mM N-acetylcysteine (Sigma-Aldrich, A9165). Media was supplemented with 5-50 ng/mL EGF (Peprotech, 315-09), 20 uM Y-27632 (Selleck Chemicals, S1049), and 1 nM DHT (Sigma Aldrich, D-073). Cells were split every 2-3 days, mechanically disrupting the cells using fire-polished pipettes each time. All cell lines were confirmed to be free of mycoplasma through regular testing using the Lonza kit (LT07-418).

### Production of virus

Lentiviruses were produced by co-transfection of 293T cells (Takara, 632180) using Lipofectamine 2000 (Invitrogen, 11668) with lentiviral backbone constructs and packaging vectors (psPAX2 and pMD2.G; Addgene, 12260 and 12259). After 2-3 days, media containing virus was collected, passed through a .45 uM filter (EMD Millipore, SE1M179M6), and concentrated (System Biosciences, LV825A-1). Virus was resuspended in organoid media containing 8 ug/mL of polybrene (EMD Millipore, TR-1003-G) and stored at -80°C (if not used immediately after harvest).

### Genetic modification of organoid lines

For lentiviral infections, established organoid lines were trypsinized (ThermoFisher, 12605010) for 5-10 min and placed in standard nonadherent cell culture plates (without Matrigel) in organoid media. The minimum amount of media needed to cover the bottom of the well was used. 8 ug/mL of polybrene (EMD Millipore, TR-1003-G) was added to the infection media improve infection efficiency. A spinfection was performed at 500 x g for 1 hour and the cells were then placed in a 37°C, 5% CO_2_ incubator overnight. Cells were collected from the plate, trypsinized briefly if any attachment has occured, and replated in 3D (Matrigel, normal conditions described above). Organoids were allowed to establish for 24 hours prior to starting antibiotic selection. Lentiviral Cre was delivered *in vitro* to recombine floxed tumor suppressor genes (*Brca2, Trp53, Pten*), and recombined cells were selected with 1 µg/mL puromycin (Invivogen, ant-pr-1).

CRISPR/Cas9 targeting of desired genes was achieved using sgRNAs in LentiCRISPRv2 containing various selection markers. Cells expressing sgRNAs targeting *Rb1* were selected with 10 µg/mL Blasticidin (Invivogen, ant-bl-1; Addgene, 98290). Post-screen, cells expressing sgRNAs against putitive resistance drivers in LentiCRISPRv2GFP (Addgene, 82416) were isolated by sorting for GFP expression 2-3 days after infection. These sgRNAs were validated sequences from the mouse Brie library (Addgene, 52963). sgRNAs were cloned into the appropriate LentiCRISPRv2 vector using established protocols (42).

### Organoid dosage response assays

Cells of interest were seeded in triplicate at 3000-3500 cells per matrigel dome per well using a 48-well non-TC treated plate (Greiner Bio-One, 677102). Drug treatments were prepared in standard mouse prostate organoid media supplemented with 5 ng/mL EGF (Peprotech, 315-09), 20 uM Y-27632 (Selleck Chemicals, S1049), and 1 nM DHT (Sigma Aldrich. D-073) at concentrations of 1 nM, 10 nM, 100 nM, 1 uM, and 10 uM. DMSO (Fisher Scientific, BP231-100) was used as control. After receiving media containing drug, cells were placed in the Incucyte S3 which was programmed to take one photo per well every six hours. % inhibition (relative to vehicle DMSO) was calculated using Microsoft Excel, and dose response curves (generated by nonlinear regression analysis) were performed using Prism 10, after 7 days of treatment.

### Positive selection CRISPR screen

Cas9 was stably expressd in the LentiCRISPR-Hygro vector, and expression was confirmed by Western bot. Doubling time for each organoid line was measured by manual counting at defined timepoints, and was ∼48h for Brca2-deficient mouse prostate organoid lines (*Brca2*^Δ*/*Δ^ *Trp53*^Δ*/*Δ^; *Brca2*^Δ*/*Δ^ *Rb1*^Δ*/*Δ^; *Brca2*^Δ*/*Δ^ *Trp53*^Δ*/*Δ^ *Pten*^Δ*/*Δ^). The Brie lentiviral library was prepared by the Gene Editing and Screening core at MSKCC (Addgene, #52963).

The pooled mouse Brie library consisted of of 4 sgRNAs per gene for 19,674 genes in the mouse genome along with 1000 nontargeting sgRNAs, where each guide is expressed in the LentiGuide-Puro vector. Brca2-deficient mouse prostate organoids were infected with the Brie sgRNA library at an MOI of 0.3 and placed under puromycin selection for four days. Cells were then either harvested (timpoint T_0_) or placed under DMSO (Fisher Scientific, BP231-100) or 100 nM olaparib (LC Laboratories) treatment for 21 days (∼10 doublings). After 21 days, cells were collected, pelleted, and resuspended in 1X PBS and submitted to the Gene Editing & Screening core at MSKCC where libraries were prepared, and the samples were sent for sequencing at the Integrated Genomics Operation at MSKCC. Screens were run in duplicate for each organoid line. Data was analyzed by the Gene Editing and Screening Core (read counts) and the Bioinformatics Core (z-scores and FDR thresholds), both at MSKCC. The guides used to individually test putative gene drivers of resistance (identified from screen) were validated sgRNA sequences from the mouse Brie library and were cloned into the LentiCRISPR-GFP (Addgene, #82416) vector to enable GFP quantification and sorting (instead of antibiotic selection).

### Immunoblotting

Cells of interest were lysed in 1X RIPA Lysis Buffer (EMD Millipore, 20-188) supplemented with phosphatase (Fisher, 52-462-51SET) and protease inhibitors (Sigma-Aldrich, 11697498001). Protein content was quantified using Pierce BCA Assay kit (Pierce, A65453) and lysates were prepared in Laemmli sample buffer (BioRad Laboratories, 1610747) supplemented with 10% 2-mercaptoethanol (BioRad Laboratories, 1610710). Samples were resolved using 4-12% Bis-Tris gel (Thermo Scientific, NP0335BOX) and blotted onto PVDF membrane (EMD Millipore, IPVH00010,) by wet transfer. Membranes were blocked in 5% milk-TBST for 1 hour before being incubated with primary antibody in 5% milk-TBST at 4 C overnight. Membranes were then rinsed in TBST and incubated with secondary antibodies in 5% milk-TBST for 1 hour. Membranes were visualized using ECL Prime kit (Cytivia, RPN2232) and autoradiography film (Genessee Scientific, 30-101L).

### GFP Competition Assay

*Brca2*^Δ*/*Δ^ GFP^+^ sgRNA-expressing cells (previously isolated by flow cytometry) were mixed with unlabelled parental cells at a ratio of 15% to 85%. These cells were plated at 2500 cells per Matrigel dome and 7500 cells per well (3 domes) in a 12-well plate. For each sgRNA tested, 2 wells for DMSO treatment and 2 wells for 100 nM olaparib treatment were plated. The proportion of GFP^+^ cells was measured every 7 days for up to 21 days.

### Immunofluorescence

Immunofluorescence for γH2AX (Abcam, ab11174) and DAPI was performed by the Molecular Cytology Core Facility (MSKCC) using established protocols and imaged on a Nikon Ti2 Eclipse microscope with a 60X oil immersion objective. γH2AX positive cells and total cells in each image were manually counted on Fiji software.

### PARP Activity Assay

Brca2-deficient organoids were treated with DMSO (vehicle) (Fisher Scientific, BP231-100), 10 nM olaparib, or 100 nM olaparib (LC Laboratories) for 7 days. Used HT PARP in vivo Pharmacodynamic ELISA Kit II (R&D Systems, 4520B-096-K) according to the manufacturers protocol for suspension cells to quantify PARP1 activity. Results were quantified on a SpectraMax M Series Multi-Mode Microplate Reader.

### DNA fiber analysis

This protocol is based on methods shared by Maria Jasin’s laboratory (MSKCC) for 2D cells and adapted for organoids grown in 3D. Mouse prostate organoid lines (*Brca2*^Δ*/*Δ^ *Rb1*^Δ*/*Δ^; *Brca2*^Δ*/*Δ^ *Trp53*^Δ*/*Δ^; *Brca2*^Δ*/*Δ^ *Trp53*^Δ*/*Δ^ *Pten*^Δ*/*Δ^) were seeded at 3,000 cells per matrigel dome and allowed to expand for 1 week. 10-20 domes are needed per condition. Organoids were dissociated with a fire-polished pipette, spun down, and resuspended in fully supplemented organoid media containing 200 µM IdU and incubated in a 37°C shaker for 20 minutes, covered from light. Cells were washed 3X with 1X PBS and resuspended in fully supplemented organoid media containing 400 µM CldU (Sigma, C6891-100MG) for 20 minutes in a 37°C shaker, covered from light. Cells were washed 3X with 1X PBS, and resuspended in fully supplemented organoid media containing DMSO or 100 µM hydroxyurea. Cells were incubated for 3 hours on a 37°C shaker, covered from light. Cells were washed with 1X PBS, trypsinized for 5 minutes in 37°C shaker, and spun down. Cells were then resuspended in 100 µL 1X PBS and placed on ice.

To place the DNA fibers, 2 uL of each sample was placed on a slide in a drop, and 7 uL of Spreading buffer was added on top and allowed to spread for 10 minutes. Cells were then slightly angled to allow the droplet to run vertically down the slide. This step should be very slow. The slides were allowed to dry for 20 minutes and then were fixed for 4 minutes in cold 3:1 methanol:glacial acetic acid solution. Slides were allowed to air dry for 5 minutes and were stored at 4°C

Slides were placed in a coplin jar containing 2N HCL for 30 minutes, covered from light. Slides were rinsed 3 times with 1X PBS and then placed in blocking buffer (5% BSA, 0.1% TritonX in 1X PBS) for 1 hour, covered from light. Slides were incubated for 1 hour, covered from light, with two primary antibodies: rat monoclonal anti-BrdU (Abcam, Ab6326, recognizes CldU) and mouse anti-BrdU (BD, 347580, recognizes IdU) by placing the antibodies at a dilution of 1:100 in 5% BSA, adding 120 uL of this mixture dropwise to each slide, and carefully place a cover slip on top. Cover slips were removed, and slides were washed 3X with 1X PBS. Slides were then incubated, covered from light, with two secondary antibodies: goat anti-mouse IgG Alexa-Fluor-488 (green) and goat anti-mouse IgG Alexa-Fluor 597 (red) by placing antibodies at a dilution of 1:250 in 5% BSA, adding 120 uL of this mixture dropwise to the slide and placing a cover slip on top. Cover slips were removed and slides were rinsed 3x with PBS and allowed to air dry, covered from light. Slides were mounted using Prolong (Invitrogen, 36930).

Slides were imaged on a Nikon Ti2 Eclipse microscope using a 60X oil immersion objective. The length of red and green tracts on a single fiber were measured in Fiji software, and a ratio of red to green color tract length, or total tract length was calculated.

### DNA extraction

Organoids were pelleted and then resuspended in 1X PBS. DNA was isolated using the Qiagen DNeasy Blood and Tissue Kit (Qiagen, 69504) according to the manufacturer’s protocol. Subcutaneous tumors were homogenized prior to DNA extraction.

### Subcutaneous mouse tumor model

NOD scid gamma mice were injected subcutaneously with 1 million cells per flank, and 5 mice were injected per organoid line, which included *Brca2*^Δ*/*Δ^ *Trp53*^Δ*/*Δ^ *Pten*^Δ*/*Δ^ organoids expressing sgRNAs against *NT1, Cdt1 g2-14, Cdt1 g2-4,* and *Cdt1 g4-2*. Tumors were measured weekly in mm^3^ and when they reached enpoint size, tjhey were harvested and flash frozen for future sample preparation.

### Sequencing of sgCdt1 clones and tumors

Clones were isolated and expanded from sgCdt1 (g2 and g4) organoids using limiting dilution. These clones were injected subcutaneously into NOD scid gamma mice as described above. Genomic DNA was extracted from organoids and tumors using the Qiagen DNeasy Blood and Tissue Kit (Qiagen, 69504) as described above. DNA samples were amplified using primers flanking the sgCdt1 cut sites and PCR products were isolated using the QIAquick Gel Extraction kit (Qiagen, 28704). Purified PCR products were sent to Integrated Genomics Operation (MSKCC) and CRISPResso2 analysis was performed to determine the editing outcomes from deep sequencing data (43). Allele fractions from pre-graft organoid lines are reported (Fig. S3) as well as from a subset of the subcutaneous tumors at the experimental endpoint (Fig. S4-S6).

#### Primary antibodies

| Antibody | Cat. no | Dilution |
| --- | --- | --- |
| β-actin | 4907S, Cell Signaling Technology | 1:10000 |
| Cas9 | 7A9-3A3, Cell Signaling Technology | 1:1000 |
| Rb1 | ab181616, Abcam | 1:1000 |
| Pten | 9559S, Cell Signaling Technology | 1:1000 |
| p53 | P53-CM5P-L, Leica | 1:1000 |
| pCHK1 (S345) | 2348S, Cell Signaling Technology | 1:1000 |
| CHK1 | 2360S, Cell Signaling Technology | 1:1000 |
| Cdc6 | 3387S, Cell Signaling Technology | 1:1000 |
| Geminin | ab246509, Abcam | 1:1000 |
| Cdt1 | PA5-42861, Invitrogen | 1:1000 |
| γH2AX | IF: ab1117, Abcam<br>WB: 2577, Cell Signaling Technology | WB 1:500 |
| H2AX | 2595S, Cell Signaling Technology | 1:1000 |

#### Drug Table

| Drug name | Cat no. |
| --- | --- |
| Olaparib | LC Laboratories |
| AZD5305 | HY-132167, MedChem Express |
| AF615 | AOB16038, AOBIOUS |
| Dimethyl sulfoxide (DMSO, Vehicle)) | BP231-100, Fisher Scientific |

## Acknowledgements

We would like to acknowledge the Histology, Molecular Cytology, Gene Editing and Screening, and Flow Cytometry cores at MSKCC. We would also like to acknowledge the integrated genomics operation (IGO) at MSKCC.

