## Supplementary figures and images for "BRCA2 reversion mutation-independent resistance to PARP inhibition in prostate cancer through loss of function perturbations in the DNA pre-replication complex"

### Figure S1

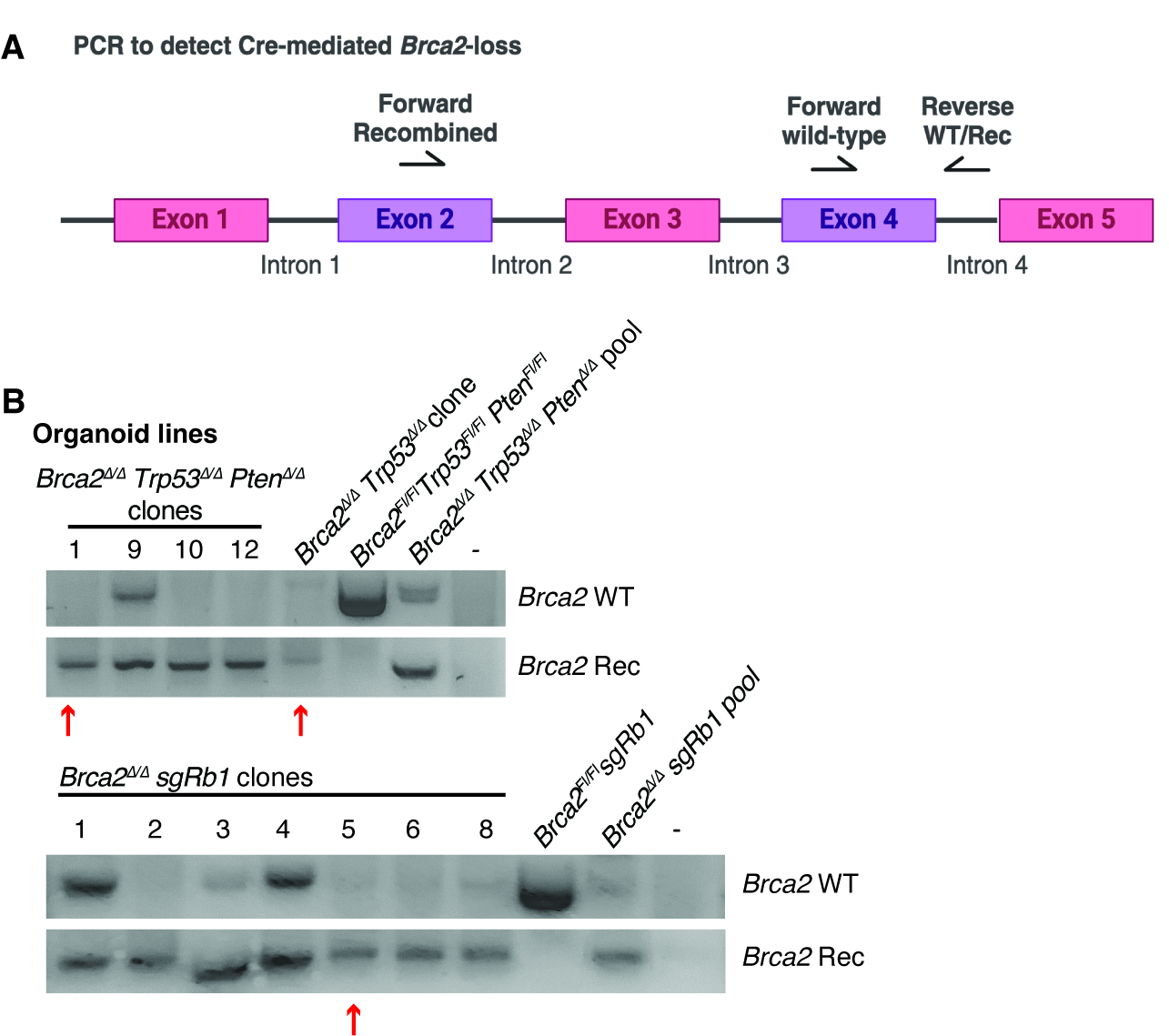

### Figure S2

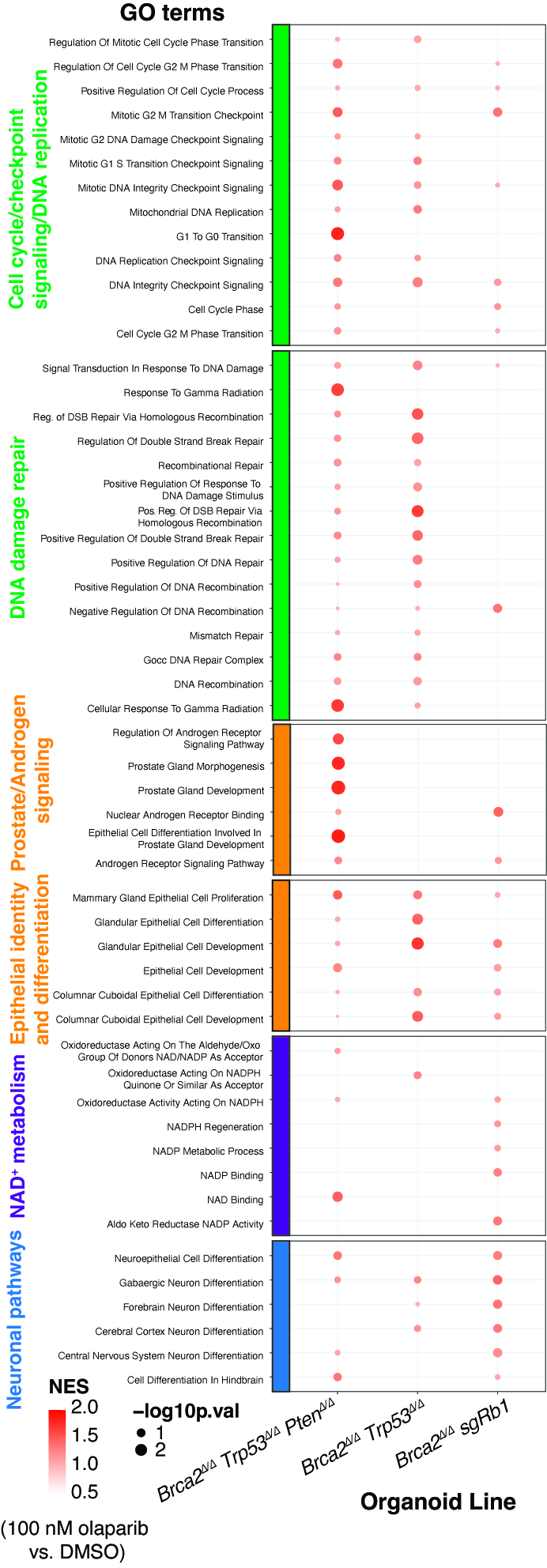

### Figure S3

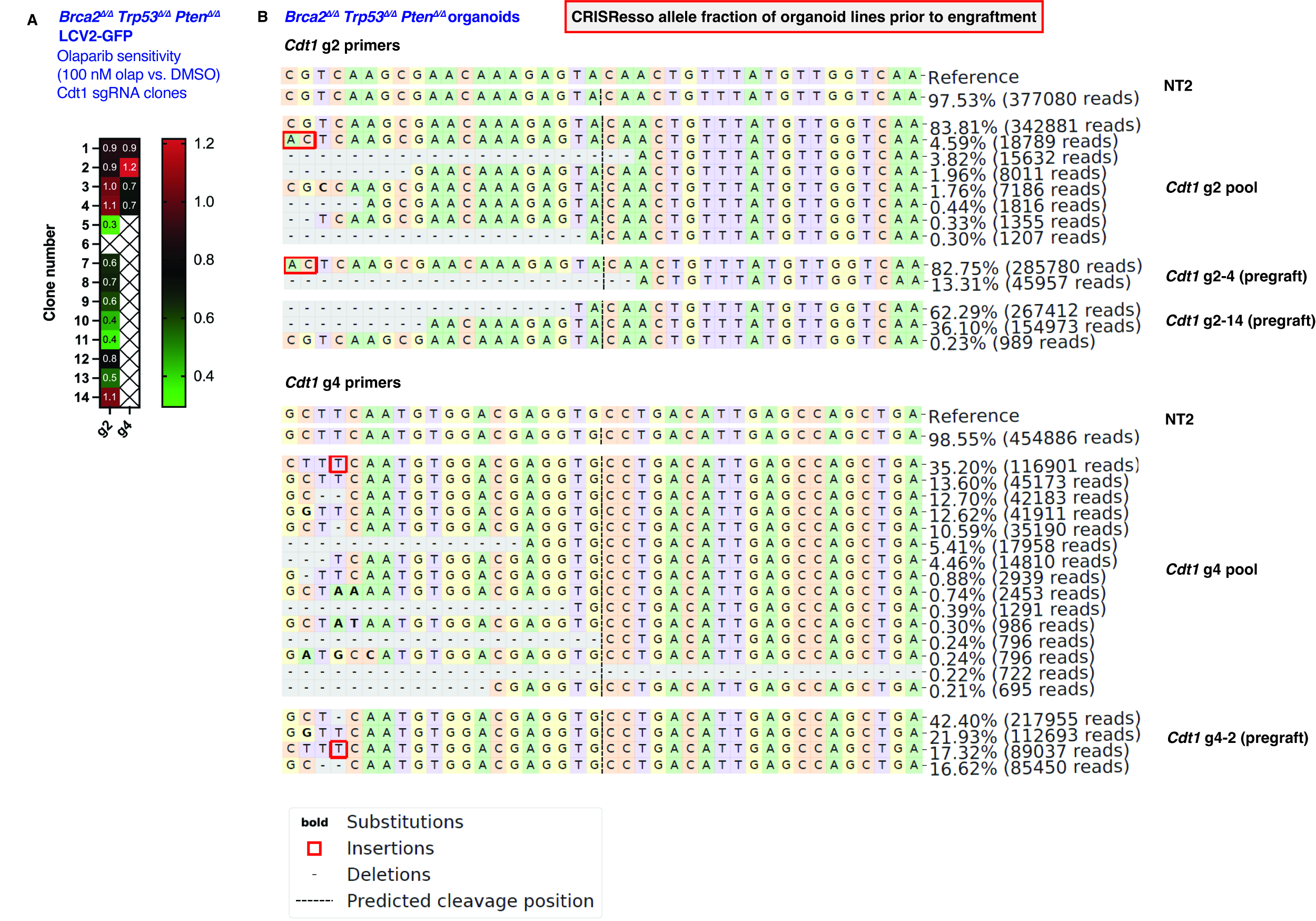

### Figure S4

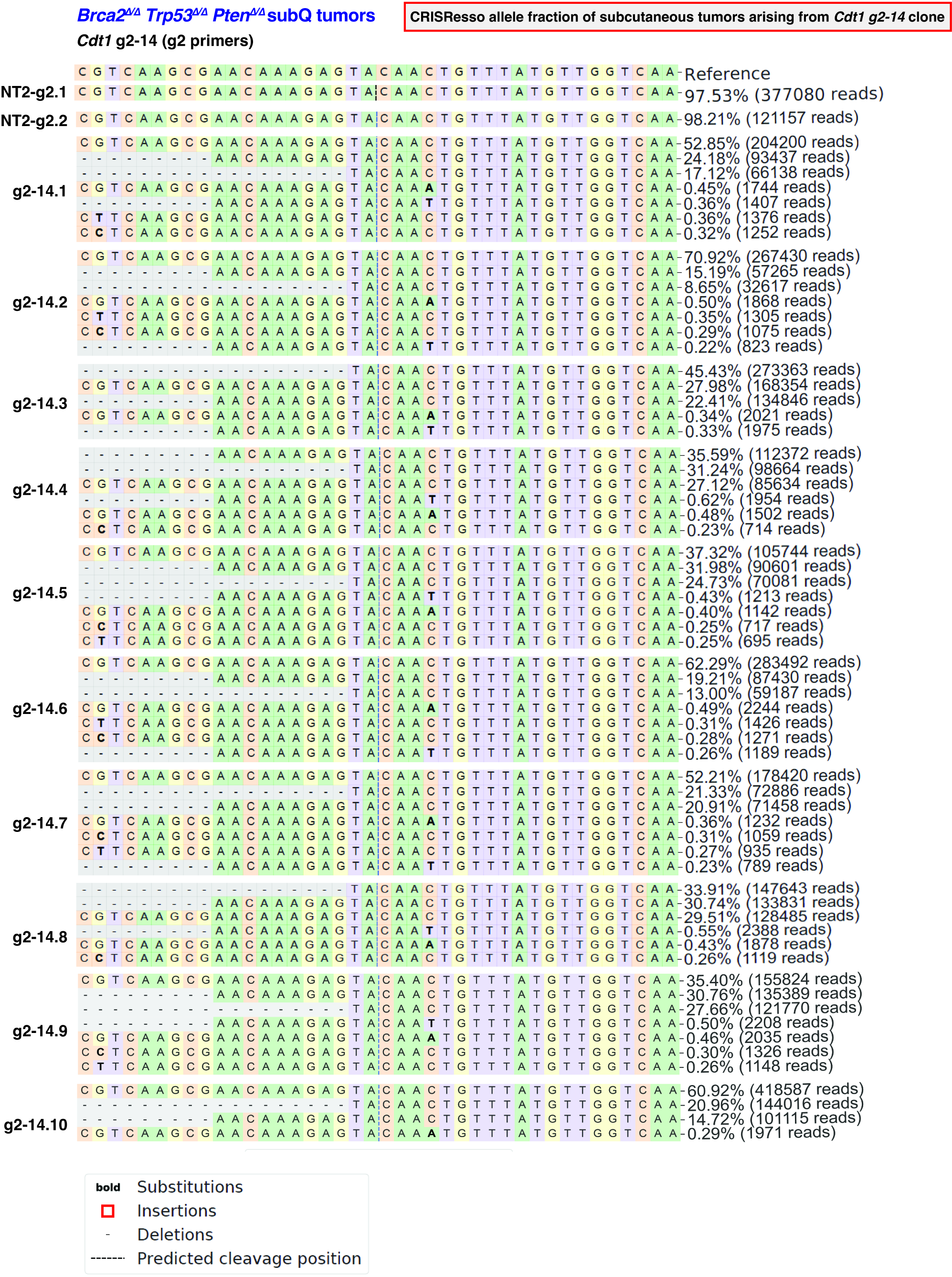

### Figure S5

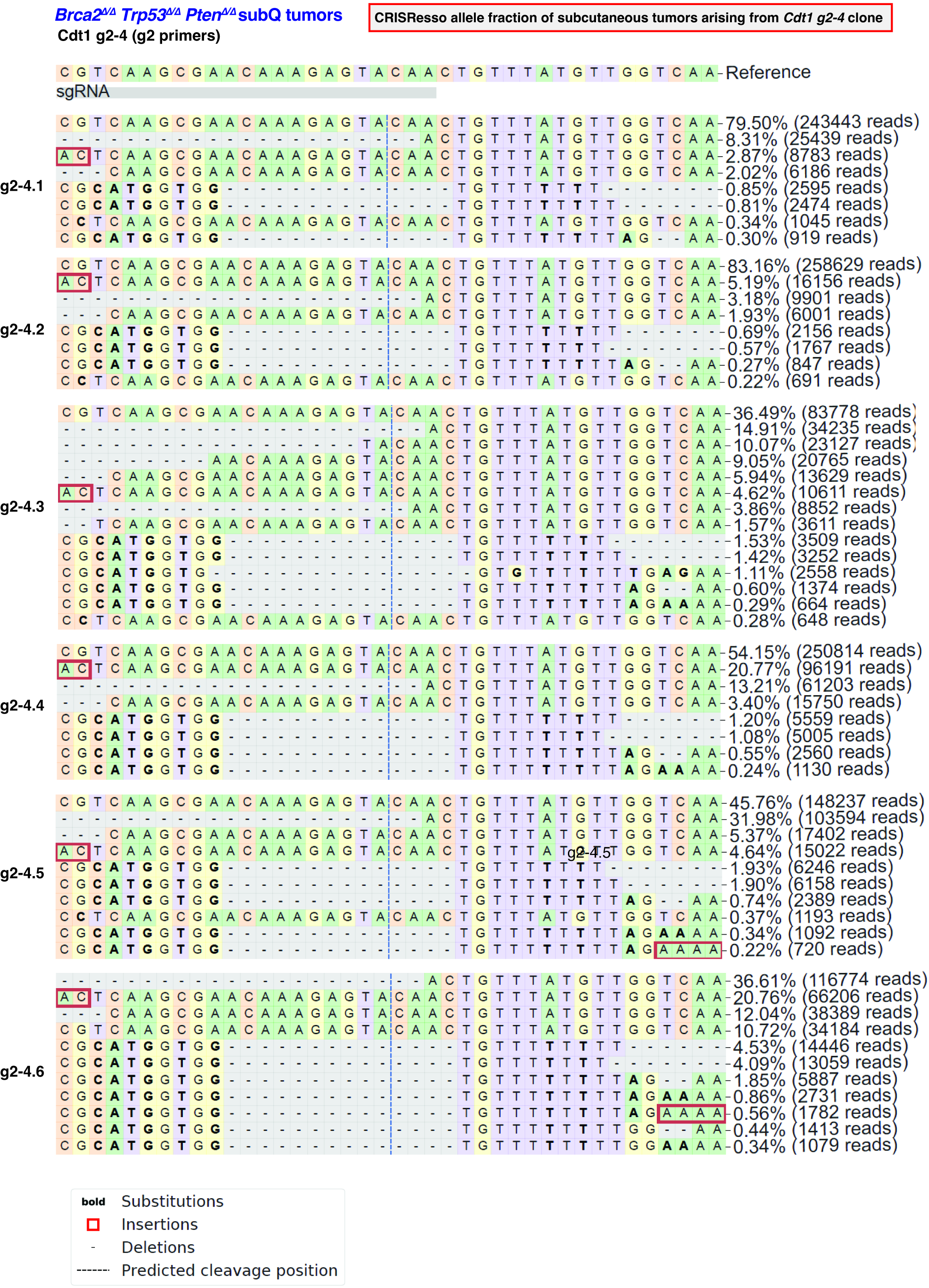

### Figure S6

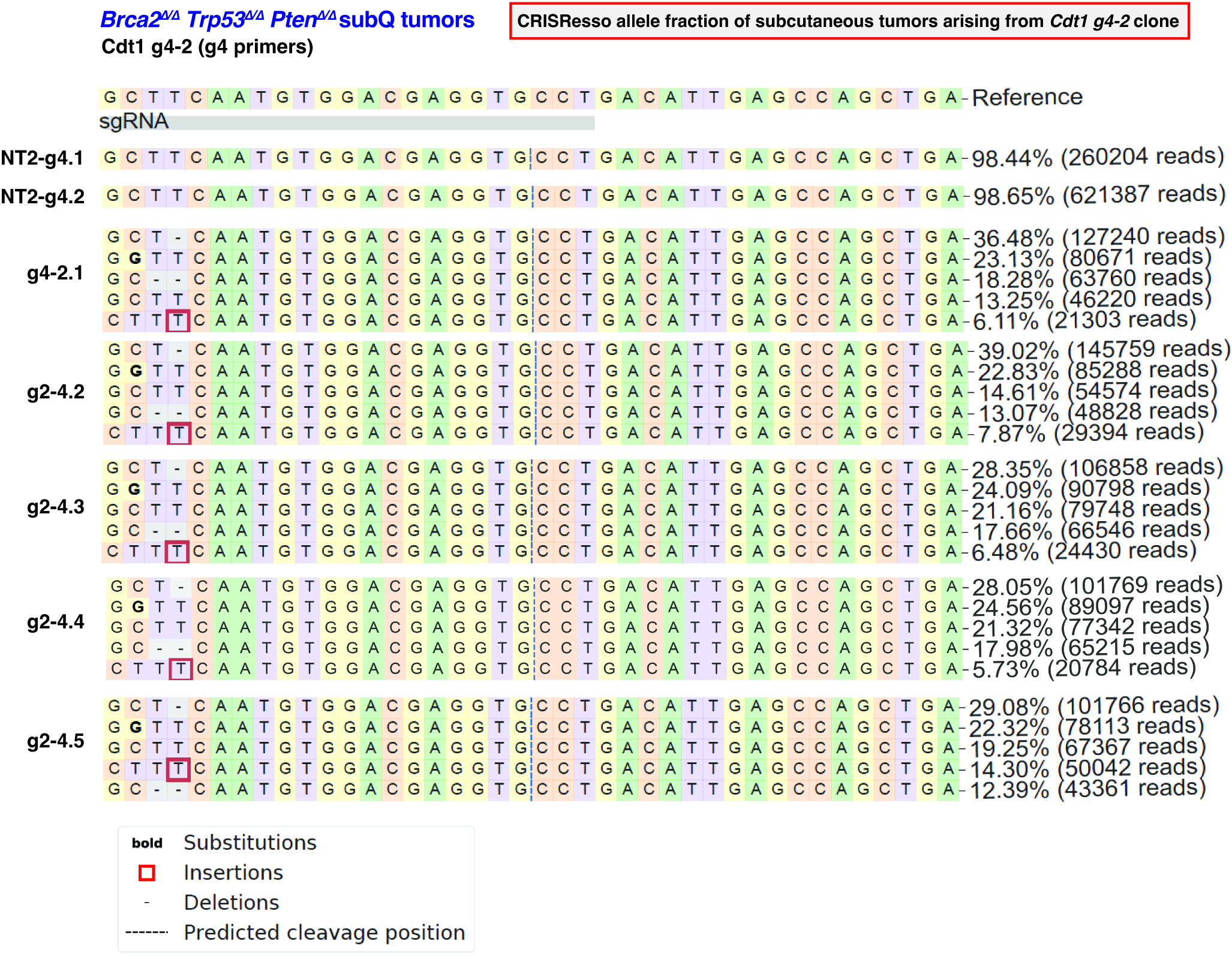

### Figure S7

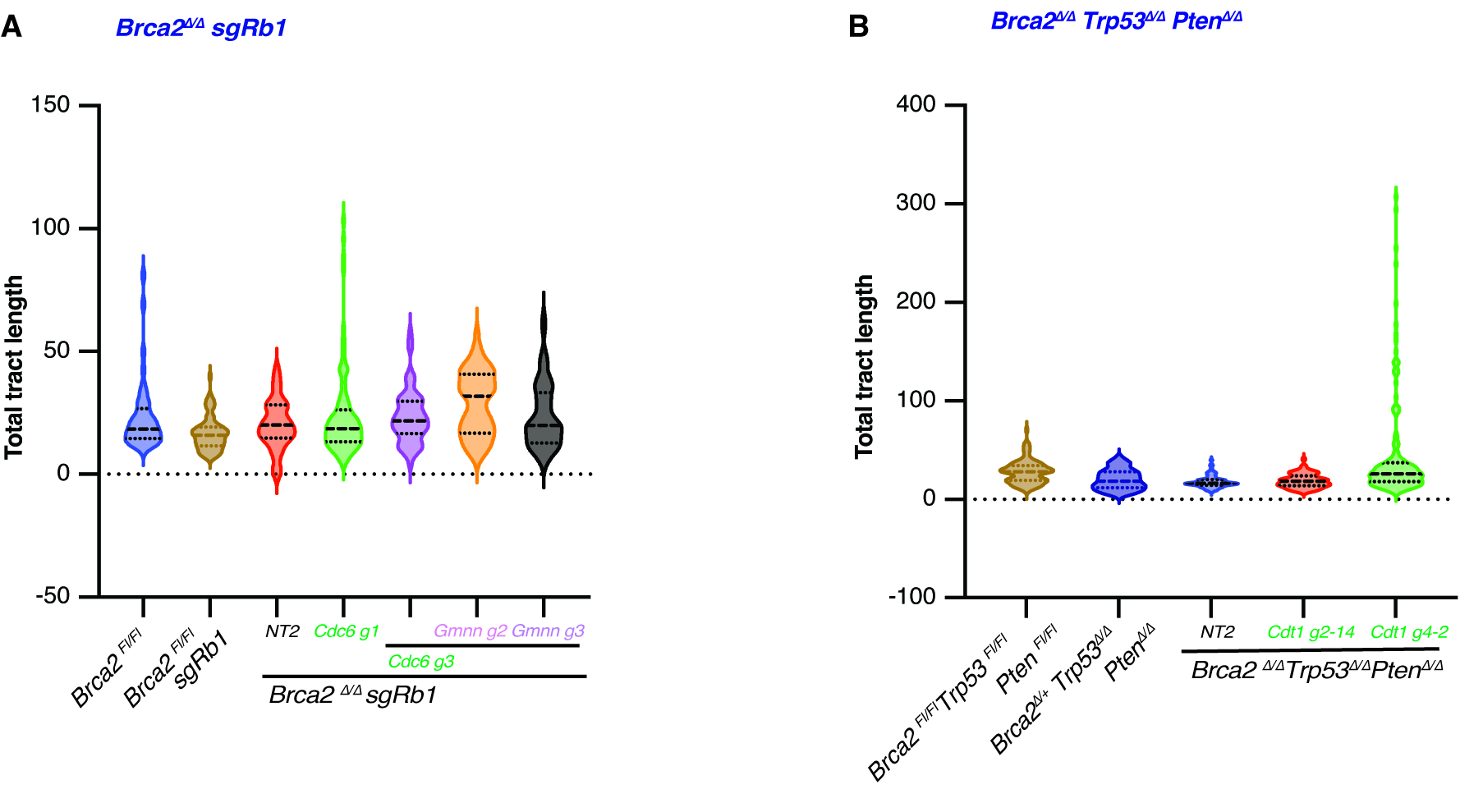
