## Supplementary Figure Legends for "BRCA2 reversion mutation-independent resistance to PARP inhibition in prostate cancer through loss of function perturbations in the DNA pre-replication complex"

**Figure S1:** Brca2 genotyping of clones following *in vitro* Cre recombination in murine prostate organoid lines. (A) Brca2 knockout was confirmed in clones using PCR primers flanking the Lox sites on the *Brca2* gene (primer locations shown on gene diagram). Amplification using the ‘Forward Recombined’ primer will only happen if recombination has occurred, and amplification using the ‘Forward wild-type’ primer will only occur if the wild-type allele is present. A knockout clone has a signal for the recombined PCR product and is negative for the wild-type PCR product. (B) Top: Brca2 genotyping of clones from *Brca2^Δ/Δ^ Trp53^Δ/Δ^ Pten^Δ/Δ^* and *Brca2^Δ/Δ^*;*Trp53^Δ/Δ^* organoids*.* Bottom: Brca2 genotyping of clones from *Brca2^Δ/Δ^sgRb1* organoids. Clones from each organoid line selected for studies are indicated by the red arrows.

**Figure S2:** Gene set enrichment analysis (GSEA) of genome-wide Brie screen in Brca2-deficient murine prostate organoid lines. GSEA was performed for all GO terms on the list of genes enriched in the genome-wide olaparib resistance screen above the 5% FDR threshold for both olaparib v. DMSO and olaparib v. T_0_ in *Brca2^Δ/Δ^ sgRb1, Brca2^Δ/Δ^ Trp53^Δ/Δ^ Pten ^Δ/Δ^, and Brca2^Δ/Δ^ Trp53^Δ/Δ^* murine prostate organoid lines. The red color gradient is the normalized enrichment score (NES), and the size of each circle represents the p-value. The scale for both parameters is indicated. Complete list of significant enrichments is reported in Supplementary Spreadsheet S2.

**Figure S3:** Characterization of *Cdt1* sgRNA clones in *Brca2^Δ/Δ^ Trp53^Δ/Δ^ Pten ^Δ/Δ/^* organoids. Two sgRNAs against *Cdt1* (guides 2 and 4) were stably expressed in *Brca2^Δ/Δ^ Trp53^Δ/Δ^ Pten^Δ/Δ^* organoids, and single clones were isolated. (A)The olaparib sensitivity of clones was evaluated by treating with 100 nM olaparib or DMSO (vehicle) for 7 days, where viability was measured on the Incucyte. Values reported are the ratio of the olaparib/DMSO signals. Red clones are resistant and green clones are sensitive on heat map. Chosen clones (*Cdt1 g2-14, g2-4, g4-2*) had sensitivity values >1. (B) DNA was amplified around g2 or g4 cleavage sites using respective primers. CRISPResso2 (1) was used to analyze sequencing data around the *sgCdt1* cut sites for g2 and g4 in organoid cultures from both guide pools and clones. The allele sequences/fraction present for g2 and g4 pools and for each chosen clone (g2-14, g2-4, g4-2) are listed. Legend for deletions, insertions, and substitutions are reported, and the predicted cleavage site is indicated by the vertical dashed line.

**Figure S4:** Sequence characterization of *Cdt1* g2-14 tumors in *Brca2^Δ/Δ^ Trp53^Δ/Δ^ Pten ^Δ/Δ/^* organoids. DNA was amplified around g2 cleavage site. CRISPResso2 (1) was used to analyze sequencing data around the *sgCdt1* cut sites for *Cdt1 g2-14* subcutaneous tumors (n=10). The allele sequences/fraction present are listed. Legend for deletions, insertions, and substitutions are reported, and the predicted cleavage site is indicated by the vertical dashed line.

**Figure S5:** Sequence characterization of *Cdt1* g2-4 tumors in *Brca2^Δ/Δ^ Trp53^Δ/Δ^ Pten ^Δ/Δ/^* organoids. DNA was amplified around g2 cleavage site. CRISPResso2 (1) was used to analyze sequencing data around the *sgCdt1* cut sites for *Cdt1 g2-4* subcutaneous tumors (n=6). The allele sequences/fraction present are listed. Legend for deletions, insertions, and substitutions are reported, and the predicted cleavage site is indicated by the vertical dashed line.

**Figure S6:** Sequence characterization of *Cdt1* g4-2 tumors in *Brca2^Δ/Δ^ Trp53^Δ/Δ^ Pten ^Δ/Δ/^* organoids. DNA was amplified around g4 cleavage site using respective primers. CRISPResso2 (1) was used to analyze sequencing data around the *sgCdt1* cut sites for *Cdt1 g4-2* subcutaneous tumors (n=5). The allele sequences/fraction present are listed. Legend for deletions, insertions, and substitutions are reported, and the predicted cleavage site is indicated by the vertical dashed line.

**Figure S7:** Tract length analysis of DNA fibers. DNA fiber analysis was performed where cells were treated with IdU (green) and CldU (red) for 20 minutes each, followed by hydroxyurea (HU) for 3h. Data is displayed as the total tract length (red + green), which is proportional to the speed of the replication fork. (A) DNA fiber tract length in *Brca2^Δ/Δ^ sgRb1* organoids stably expressing the indicated sgRNAs *(sgNT1, sgNT2, Cdc6 g1, Cdc6 g3, Gmnn g2, Gmnn g3)* in LentiCRISPR-GFP. *Brca2^Fl/Fl^ and Brca2^Fl/Fl^sgRb1* organoids were included as controls. (B) DNA fiber tract length in *Brca2^Δ/Δ^ Trp53^Δ/Δ^ Pten^Δ/Δ^* organoids expressing the indicated sgRNAs (*sgNT1, Cdt1 g2-14, Cdt1 g4-2*) in LentiCRISPR-GFP. *Brca2^Fl/Fl^ Trp53^Fl/Fl^ Pten^Fl/Fl,^* and *Brca2^Δ/+^ Trp53^Δ/Δ^ Pten^Δ/Δ^* organoids were included as controls.
